# High-resolution 2D Solid-State NMR provides insights into Nontuberculous Mycobacteria

**DOI:** 10.1101/2024.05.28.596255

**Authors:** Chang-Hyeock Byeon, Kasper Holst Hansen, William DePas, Ümit Akbey

## Abstract

We present a high-resolution magic-angle spinning (MAS) solid-state NMR (ssNMR) study to characterize native nontuberculous mycobacteria (NTM). We studied two different NTM strains, *Mycobacterium smegmatis*, a model, non-pathogenic strain, and *Mycobacterium abscessus*, an emerging and important human pathogen. Native hydrated NTM samples were studied at natural abundance without isotope-labelling and any chemical or physical modification. We utilized 1D ^13^C and 2D ^1^H-^13^C ssNMR spectra and peak deconvolution to identify NTM cell-wall chemical sites. More than ∼100 distinct ^13^C signals were identified in the ssNMR spectra. The signals originating from both the flexible and rigid fractions of the native bacteria samples were selectively analyzed by utilizing either CP or INEPT based ^13^C ssNMR spectra. CP buildup curves provide insights into the dynamical similarity of the cell-wall components for NTM strains. Signals from peptidoglycan, arabinogalactan and mycolic acid were identified. We also provide tentative assignments for ∼30 polysaccharides by using well resolved ^1^H/^13^C chemical shifts from the 2D INEPT-based ^1^H-^13^C ssNMR spectrum. As an orthogonal way of characterizing the bacteria, electron microscopy (EM) was used to provide spatial characterization. ssNMR and EM data suggest that *M. abscessus* cell-wall is composed of a smaller peptidoglycan layer which is more flexible compared to *M. smegmatis*, which may be related to its higher pathogenicity. Here in this work, we used high-resolution 2D ssNMR first time to characterize native NTM strains and identified chemical sites. These results will aid the development of structure-based approaches to combat NTM infections.

## 1. Introduction

Antimicrobial resistance (AMR) can, render conventional infection treatments ineffective and therefore poses a significant global health threat.^1^ This escalating crisis not only amplifies the complexity of treating common infections but also exacerbates the burden on healthcare systems worldwide. Chronic infections further compound this issue, as they require prolonged and often intensified antimicrobial therapies, placing a higher strain on patients and healthcare systems and increasing the risk of resistance development. Within this landscape, nontuberculous mycobacteria (NTM) stand out as opportunistic pathogens capable of causing a spectrum of recalcitrant infections.^2^ The intrinsic resistance of NTM to many conventional antibiotics underscores the urgency of addressing AMR comprehensively, highlighting the imperative for innovative therapeutic strategies and robust antimicrobial stewardship efforts.

Nontuberculous mycobacterial infections have outnumbered *Mycobacterium tuberculosis* infections in the United States for over a decade,^3^ and the incidence and prevalence of NTM infections continues to increase.^4^ NTM can cause skin and soft tissue infections, but they more frequently cause pulmonary disease in patients with a compromised immune system or underlying lung disorders such as Cystic Fibrosis (CF) and non-CF bronchiectasis.^5-7^ Indeed, NTM affects up to 32% of people with CF over the age of 40, and 8-10% of the 350,000 to 500,000 non-CF bronchiectasis patients in the US are culture-positive for NTM.^7-10^ *Mycobacterium abscessus* is one of the major NTM pathogens, is particularly difficult to treat with conventional antibiotics, and is associated with more severe clinical symptoms compared to other NTM species and to common Gram-negative pathogens.^9, 11-13^ Treatment failure rates for *M. abscessus* pulmonary infections can approach 75%, and lung resections are sometimes employed for localized *M. abscessus* infections.^14, 15^

Mycobacteria and related members of the *Mycobacteriales* order feature a cell envelope structure that is distinct from typical Gram-negative or Gram-positive bacteria.^16^ The mycobacterial surface is composed of an inner membrane, a periplasmic space, and peptidoglycan (PG). Then, a unique arabinogalactan (AG) layer links the peptidoglycan to an outer layer referred to as the mycomembrane (MM).^16-18^ The inner leaflet of the mycomembrane is composed of long chain mycolic acids (MA) that are covalently linked to the arabinogalactan layer. The outer leaflet and capsule layer include a variety of free mycolic acids, proteins, polysaccharides, and lipoglycans.^16, 19^ The cell-wall of *Mycobacteriales* is a direct target for antimicrobial drugs.^20^ The composition of the mycomembrane impacts host-pathogen interactions, biofilm formation, and antibiotic tolerance in both NTM and *M. tuberculosis*.^21-25^ An improved understanding of the mycobacterial envelope is pivotal for developing novel therapeutic interventions and diagnostic strategies to combat NTM infections, particularly in the context of rising antimicrobial resistance.

Magic-angle spinning (MAS) solid-state NMR (ssNMR) can provide high-resolution structural and dynamics information.^26-30^ Compared to other techniques, ssNMR is extremely powerful and can be utilized to study native systems at physiological conditions even without purification as a whole (whole cell ssNMR),^31^ such as bacteria, biofilms and others. In recent years ssNMR spectroscopy has been widely used to characterize such systems on bacterial, plant, and fungal cell walls.^32-34^ For bacterial cell wall chemical characterization, solution NMR and chemical decomposition coupled with mass-spectrometry methods have been used.^35, 36^ These two methods rely on the solubility of the chemicals at the studied systems, and quantification is usually limited due to the difficult to solubilize/extract insoluble species. In this respect, ssNMR has a great advantage and can study the systems as prepared as a whole and intact, e.g. both soluble and insoluble fractions at once.^37, 38^

By utilizing different polarization schemes, such as CP versus INEPT, ssNMR could differentiate signals from flexible versus rigid signals from corresponding chemical species. We demonstrated this on natural-abundance native *Pseudomonas fluorescens* bacterial colony biofilms recently.^39^ To date, predominantly 1D ssNMR experiments have been utilized to obtain structural information on bacterial, fungi, plant systems.^32, 33, 40^ Despite severe signal overlap in 1D ssNMR spectroscopy, many structural details have been extracted from these and correlated to the nature of the samples. Multidimensional (nD) ssNMR is the way to reduce the redundancy of the overlapping signals by adding an extra chemical shift dimension, proton or carbon. Despite very powerful, this method usually require longer experiment times or isotope-labelling (^13^C and/or ^15^N), which is not possible for studying natural extracts and can only be done in lab grown systems. When the system of interest is from e.g. patients, or isotope-labeling is not possible, natural-abundance samples have to be utilized which limits the ssNMR experimental approaches to either 1D or simpler 2D experiments utilizing proton as the second dimension. We recently showed such a 2D ^1^H-^13^C correlation as approach as a powerful method to investigate *Pseudomonas* biofilm samples. Unfortunately, natural abundance 2D ^13^C-^13^C spectra is not feasible with conventional ssNMR approaches. Hyperpolarized nD ssNMR can overcome this sensitivity limitation and has been applied to different natural abundance systems successfully, with a caveat of lower resolution due to cryogenic experimental temperatures.^34, 41-43^ In summary, a small number of nD ssNMR application on complex natural systems have been demonstrated so far, such as on cell-walls of ^13^C-labeled *S. aureus*,^44 13^C-labeled *E. coli*-*P. aureginosa* cell-wall,^45, 46^ and ^13^C-labeled *B. subtilis* cell-wall.^43^ Moreover, there are a sizeable amount of solution NMR studies on cell-wall materials.^36, 47, 48^

Previously 1D ^13^C CPMAS ssNMR was utilized by Cegelski and coworkers to obtain structural information about the cell envelope of NTM.^49^ They analyzed the 1D ssNMR spectra of a lyophilized dry *M. smegmatis* whole cell sample, chemically extracted cell-wall components, and commercial mycolic acid and arabinogalactan materials. Their ssNMR spectra of the extracted cell-wall material (PG+AG+MA) and whole-cell were different, highlighting the importance of studying native untreated bacteria samples. Moreover, this work relied on 1D ssNMR spectroscopy, and despite being powerful it poses challenges in signal identification due to lower-resolution and signal overlap. In addition to the NMR work, fractionation analysis and cryo-electron tomography has been used to characterize the cell envelope of NTM. ^16-18, 22^

Here, we present a MAS ssNMR study to obtain structural and dynamics information on native NTM towards a better understanding of molecular structure. We characterized hydrated intact native whole-cell NTM that were grown in liquid medium without chemical extraction/fractionation of individual components. We then packed the bacteria into the ssNMR rotors in their natural hydrated whole cell form without any further drying or processing. We utilize conventional room-temperature carbon or proton-detected 1D and 2D MAS ssNMR to characterize and quantify both native nonpathogenic *M. smegmatis* and pathogenic *M. abscessus*. We obtained high-resolution structural as well as qualitative dynamics information. This is the first high-resolution 2D ssNMR study on NTM that identifies chemical sites without signal overlap and ∼30 unique polysaccharide sites. Utilizing ssNMR spectroscopy and electron microscopy on NTM represents progress in our understanding of the molecular basis of NTM cell envelope architecture. By unraveling these structural properties of the NTM cell envelope, we contribute to future development of innovative strategies for combating NTM-related diseases.

## 2. Methods

### Bacteria strains and preparation

*Mycobacterium smegmatis* MC^2^155 and *Mycobacterium abscessus* smooth clinical isolate 0253a, a pulmonary infection isolate from a person with Cystic Fibrosis,^50^ were grown in liquid TYEM (10 g tryptone and 5 g yeast extract per liter with 2 mM MgSO_4_) for 48 hours (*M. smegmatis*) or 72 hours (*M. abscessus*) at 37°C, shaking at 250 rpm. At these timepoints and in these conditions, aggregates had fully dispersed into planktonic cells.^50^ Cultures were centrifuged for 10 minutes at 7000 x g at 4°C, and the supernatant was removed. Fixed samples were resuspended in PBS + 4% paraformaldehyde and gently rocked at room temperature for 3 hours. Fixed samples were centrifuged again, the supernatant was removed, and the pellet was kept at 4 °C until the ssNMR measurement. Unfixed *M. smegmatis* sample pellets were put on ice and measured immediately by ssNMR.

### NMR Spectroscopy

All MAS ssNMR experiments were performed at 750 MHz Bruker Avance 3 spectrometer equipped with a low-temperature triple-resonance 3.2 mm probe. We utilized thin-walled 3.2 mm rotors for the measurements. Total amount of material in the rotor was ∼50 mg. We used a benchtop centrifuge to transfer the pellets into the rotor, by using pipets attached directly onto the rotor and then centrifuged. We employed 10 kHz MAS for 1D experiments and 20 kHz MAS for 2D experiments. The set temperature was 275 K which corresponds to a sample temperature of around ambient temperature. 3.3 μs and 5 μs pulses were used for ^1^H and ^13^C. For the CP experiments, 1 ms contact time was used with a 70-100% ramp on the proton channel. ∼90 kHz proton dipolar decoupling was applied for all the spectra recorded. The ^1^H chemical shifts were referenced directly to 0 ppm by using DSS as an internal standard added to the NMR samples, and the ^13^C chemical shifts were indirectly referenced by using the ^1^H frequency.^51^

For both NTM samples, the 1D ^13^C INEPT, cross-polarization (CP) and direct polarization (DP) spectra were recorded with 32k, 32k and 7k scans, respectively, to allow for good SNT for analysis in ∼9 hours. 1 second of recycle delays were used for CP and INEPT, and 5 seconds for DP experiment. The spectral fitting and peak deconvolution were performed by using the ssNake program package.^52^ The 1D spectra were processed with gaussian broadening window function (with 35/100 Hz at GB=0.02 in Topspin).

The carbon-detected 2D ^1^H-^13^C (HC) INEPT ssNMR spectra were recorded with 1k transients in ∼22 hours with 20 kHz MAS. 1 s recycle delay was used and a total of 78 indirect dimension data points were recorded with an increment of 100 μs. The proton-detected 2D ^1^H-^13^C (hCH) INEPT ssNMR spectra were recorded with 128 transients in ∼3 hours with 20 kHz MAS. 1 s recycle delay was used and a total of 90 indirect dimension data points were recorded with an increment of 40 μs. The proton-detected 2D ^1^H-^13^C (hCH) CP ssNMR spectra were recorded with 196 transients in ∼9 hours with 20 kHz MAS. 1 s recycle delay was used and a total of 160 indirect dimension data points were recorded with an increment of 28 μs. The 2D spectrum was processed with gaussian broadening (with 35 Hz at GB=0.025 in Topspin) for direct dimension and by a mixed sine squared for indirect dimension (SSB=3 in Topspin).

We utilized the ^1^H and ^13^C chemical shift values given in the CCMRD database for identification of polysaccharide chemical sites in our ssNMR spectra. We only judge the presence of a certain polysaccharide species with at least three chemical shift matches identified from each specific chemical candidate.

## 3. Results

### 3.1. Structural features of native NTM by electron microscopy and ssNMR

The chemistry of cell wall components contributes to the unique properties of NTM, including their resistance to antibiotics, ability to persist in the environment, and capacity to cause chronic infections, **Figure 1**. Understanding the biochemical composition of the NTM cell envelope is crucial for developing targeted therapies to combat NTM infections effectively. Unlike typical bacterial cell walls, NTM possess a complex outer layer composed primarily of peptidoglycan, arabinogalactan, and uniquely mycolic acids long-chain fatty acids that form a lipid-rich barrier. Different structural elements forming the NTM cell-wall have very different dynamics, flexible versus rigid, and poorly understood.^53^

**Figure 1.**
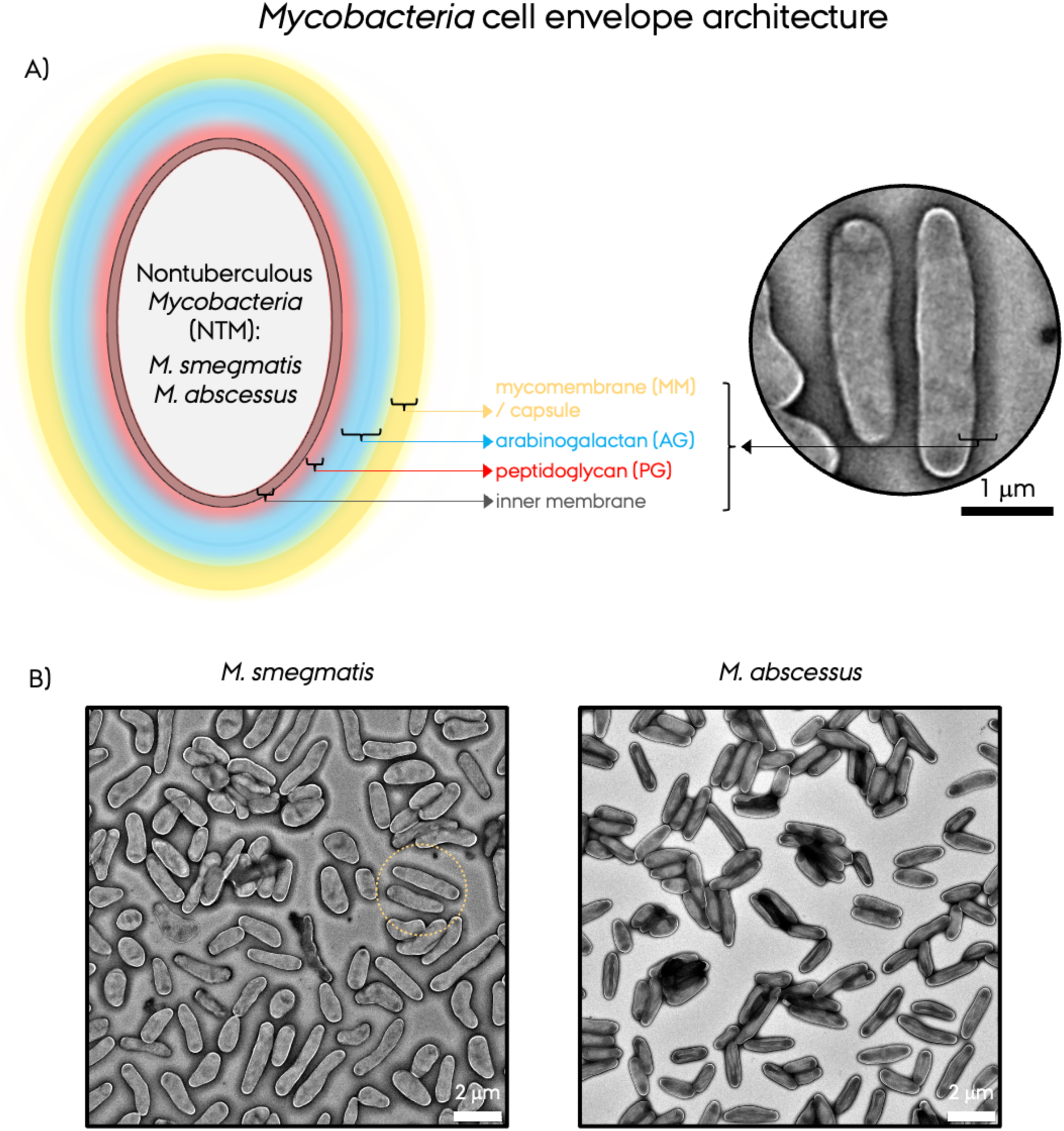
**A)** A cartoon representation of the NTM cell envelope architecture and its components, such as inner membrane, peptidoglycan (PG), arabinogalactan (AG), and mycomembrane (MM)/capsule. The zoomed out image is taken from the fixed *M. smegmatis* EM micrograph marked as yellow dashed-circle. **B)** Negative staining EM micrographs of planktonic *M. smegmatis* and *M. abscessus*.

Mycolic acids, which are long-chain fatty acids characterized by their high degree of lipid content and branching. Mycolic acids form a hydrophobic envelope which provides NTM with impermeability to many hydrophilic compounds, including antibiotics and detergents. This lipid-rich barrier is essential for NTM survival within host tissues and interact with host cells and the immune system. The cell wall lipids, such as glycolipids and phospholipids, contribute to the NTM architecture. Glycolipids, including trehalose dimycolate and glycopeptidolipids are functionally important. Embedded within the mycolic-acid-rich mycomembrane are various proteins, glycolipids, and polysaccharides (Fig. 1A).^16^ Mycolic acid and lipids give ^13^C chemical shifts at ∼32.5 and ∼132 ppm due to methylene and alkene signals, respectively.^49^

Peptidoglycan is another component of the NTM cell envelope although present in smaller amounts compared to *M. tuberculosis*.^54^ Peptidoglycan provides structural support to the cell envelope and is crucial for maintaining cell shape and integrity. It consists of alternating N-acetylglucosamine (GlcNAc) and N-acetylmuramic acid (MurNAc) residues cross-linked by peptide bridges that are composed of aminoacids, forming a mesh-like structure. Proteins embedded within the NTM cell wall serve various functions, including cell adhesion, nutrient transport, and signal transduction. These proteins interact with host cells and mediating NTM’s interactions. PG gives ^13^C chemical shifts at a wide range of ∼10-180 ppm. In contrast, the arabinogalactan is composed of galactose, arabinose and rhamnose with ^13^C shifts appear at ∼50-110 ppm. The arabinogalactans are chemically linked to peptidoglycans and mycolic acids, supplying structural rigidity.

EM and NMR methods can provide different types of information about these bacteria, as EM can provide high-resolution images of the overall structure of a sample, whereas ssNMR can provide detailed information about the chemical composition and molecular interactions within that structure. For the first level of characterization of our two distinct NTM strains, *M. smegmatis and M. abscessus*, we utilized negative-staining EM, **Figure 1**. The elongated rod-shaped cells are visible at the EM micrographs. The zoomed out EM images indicate the cell envelope as a thick dark layer that expands outside the bacteria, **Figure 2A**. The planktonic *M. smegmatis and M. abscessus* appear similar in shape. The cell envelope thickness of *M. abscessus* appears to be slightly thinner than *M. smegmatis*. The level of structural details at these negative EM images are useful but limited in understanding the chemical composition, so we applied ssNMR to get more detailed structural information.

**Figure 2.**
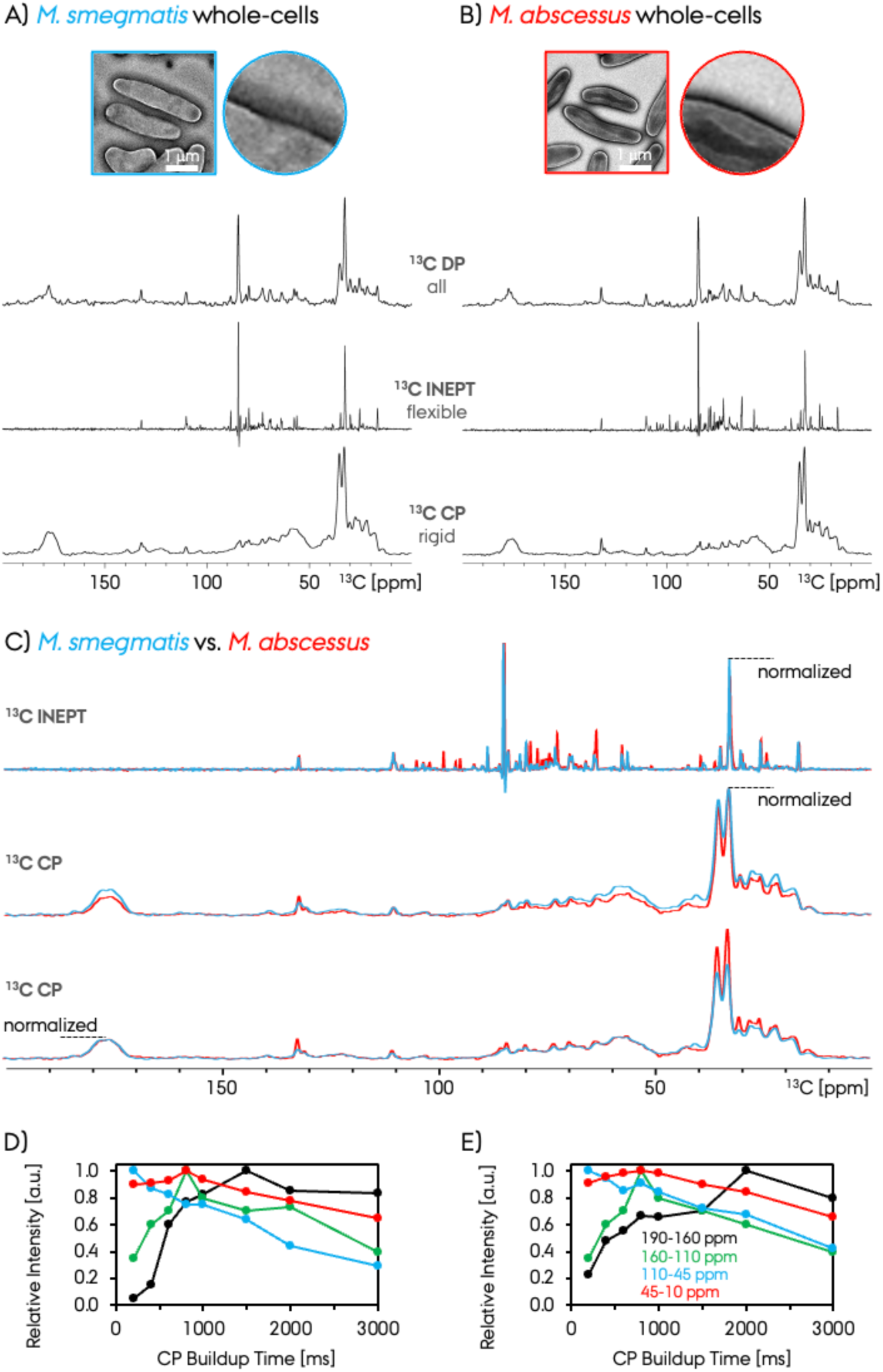
**A,B)** 1D ^13^C ssNMR spectra recorded on wet and dry biofilm preparations. CP and INEPT polarization transfer schemes were used to record the ^13^C ssNMR spectra. **C)** The comparison of *M. smegmatis* and *M. abscessus* ^13^C INEPT, CP and DP spectra. The different points for intensity normalization are shown with dashed lines. The CP buildup for different signals in the 1D ^13^C CPMAS spectra (given with the color-coded curves as a functional of contact-time) are shown for **D)** *M. smegmatis* and **E)** *M. abscessus* NTM samples. Each CP buildup curve was normalized to the maximum intensity within that same dataset and different curve intensities are not directly comparable. The four different chemical shift ranges for the integration of signals are given with color coding in terms of integration ppm scale at the ^13^C CP ssNMR spectra.

The ssNMR spectra shown in **Figure 2** demonstrates the feasibility of detecting sufficient ^13^C ssNMR signals at natural abundance from native NTM with sufficient sensitivity within few hours. We utilized the conventional ssNMR at ambient temperature and moderate MAS frequencies (10 or 20 kHz). The samples were packed into a ssNMR rotor as close to the *in vivo* condition by gentle centrifugation as wet pellets without any further treatment. NTM was fixed post-harvesting with paraformaldehyde in PBS buffer to allow stability and long term durability. As a result, we were able to record ssNMR spectra at close to native conditions, **Figures 1** and **2**. The fixation of the *M. smegmatis* shows only small changes in the mycolic acid region in the ssNMR spectra compared to the unfixed sample, as shown with dashed-circles in **Figure 3**. The rest and majority of the spectra were not affected. Below, 1D and 2D ssNMR results are discussed in detail and correlated to NTM cell envelope architecture.

**Figure 3.**
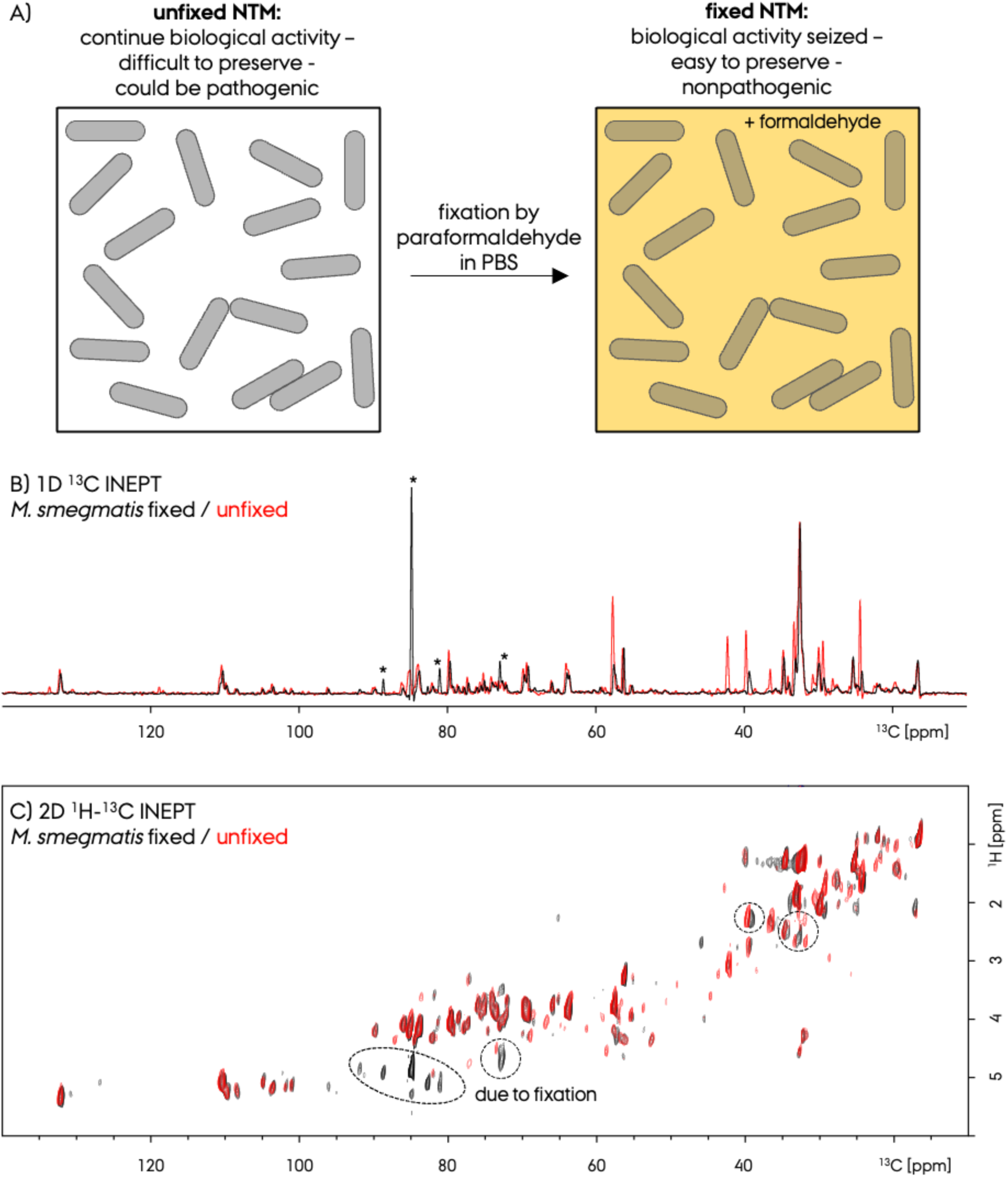
**A)** Schematic representation of fixed and unfixed planktonic *M. smegmatis*. **B)** Comparison of 1D ^13^C INEPT ssNMR spectra of fixed and unfixed *M. smegmatis*. The peaks marked with asterisk are due to paraformaldehyde in PBS buffer that is used for fixation. **C)** Comparison of 2D ^1^H-^13^C INEPT ssNMR spectra of fixed and unfixed *M. smegmatis*.

### 3.2. 1D ^13^C ssNMR for native NTM characterization

High-sensitivity 1D ^13^C ssNMR spectra were recorded with ∼50 mg of native NTM samples of *M. smegmatis* and *M. abscessus* in ∼9 hours, **Figure 2A,B**. These spectra were recorded with CP and INEPT which selectively report on the rigid versus flexible biofilm components, respectively. Additionally, we recorded direct polarization (DP) spectra to obtain rigid and flexible signals at once which is a linear combination of the CP and INEPT spectra. We previously quantified the rigid versus flexible parts of a *Pseudomonas* biofilm sample,^39^ moreover, others reported on similar characterization of bacterial cell walls.^46^ For *M. smegmatis* and *M. abscessus*, this comparison has not been previously shown. The INEPT and CP spectra recorded at room temperature for both samples have significantly different resolution, due to the very different dynamics of the signals observed. The average ^13^C resonance linewidth is ∼50 Hz (∼0.3 ppm) and ∼440 Hz (∼2.4 ppm) for the deconvoluted peaks in the 1D INEPT and CP spectra, respectively, **Table 1**. The ratio of the CP/INEPT ^13^C signal integrals are noticeably different for spectra recorded under similar conditions for the *M. smegmatis* and *M. abscessus*, 0.75 versus 0.40, respectively. This indicates that both of the native NTM strains consist of less rigid chemical species relative to the flexible species. Remarkably, the relative rigid fraction of *M. smegmatis* is much larger than *M. abscessus* by almost two-fold, which is also seen in the spectral intensity differences in **Figure 2C**.

**Table 1:** The list of identified chemical sites in the 1D ^13^C INEPT and CP ssNMR spectra for the native fixated *M. smegmatis* sample, as shown in Figure 4. Chemical shifts in terms of ppm, linewidths in terms of Hz/ppm, and relative abundance percentage integral ratios are given. The normalization of the abundance percentage was done for each chemical site by dividing its integral to the total integral sum within CP or INEPT spectrum separately. The intense paraformaldehyde signal is omitted for relative abundance calculation in the INEPT spectrum. The average ^13^C resonance linewidth is ∼50 Hz (∼0.3 ppm) and ∼440 Hz (∼2.4 ppm) for the deconvoluted peaks in the 1D INEPT and CP spectra, respectively.

| *M. smegmatis* <sup>13</sup>C INEPT ssNMR |  |  |  |  | *M. smegmatis* <sup>13</sup>C CP ssNMR |  |  |  |
| --- | --- | --- | --- | --- | --- | --- | --- | --- |
| ppm | ppm | Hz | % |  | ppm | ppm | Hz | % |
| 16.6 | 0.2 | 42 | 5.2 |  | 14.2 | 1.7 | 316 | 0.5 |
| 18.4 | 0.2 | 43 | 0.2 |  | 18.2 | 3.1 | 576 | 4.0 |
| 19.3 | 0.2 | 39 | 0.4 |  | 21.9 | 3.1 | 586 | 4.5 |
| 19.7 | 0.3 | 49 | 0.3 |  | 23.3 | 3.3 | 619 | 3.1 |
| 20.3 | 0.3 | 58 | 0.4 |  | 25.6 | 1.8 | 338 | 2.9 |
| 21.1 | 0.4 | 84 | 0.2 |  | 27.5 | 2.1 | 400 | 4.7 |
| 21.8 | 0.3 | 55 | 1.0 |  | 30.1 | 2.1 | 398 | 4.1 |
| 22.1 | 0.3 | 49 | 0.5 |  | 32.1 | 1.5 | 280 | 1.0 |
| 23.8 | 0.3 | 55 | 1.5 |  | 32.9 | 1.8 | 344 | 12.3 |
| 24.6 | 0.5 | 87 | 1.4 |  | 33.8 | 1.3 | 247 | 0.2 |
| 25.3 | 0.2 | 35 | 3.4 |  | 35.3 | 2.1 | 392 | 13.0 |
| 27.6 | 0.3 | 52 | 0.5 |  | 40.3 | 2.5 | 469 | 2.8 |
| 29.2 | 0.2 | 31 | 0.7 |  | 42.7 | 2.5 | 469 | 1.8 |
| 29.9 | 0.3 | 55 | 4.2 |  | 45.5 | 2.5 | 469 | 1.0 |
| 32.5 | 0.3 | 53 | 21.8 |  | 50.5 | 3.0 | 563 | 1.3 |
| 34.7 | 0.3 | 51 | 4.3 |  | 52.9 | 3.0 | 563 | 2.5 |
| 38.2 | 0.2 | 45 | 0.8 |  | 56.0 | 3.6 | 680 | 4.5 |
| 38.6 | 0.2 | 44 | 1.1 |  | 57.9 | 3.5 | 657 | 1.6 |
| 42.1 | 0.3 | 55 | 0.7 |  | 59.4 | 2.9 | 551 | 2.7 |
| 55.0 | 0.2 | 34 | 0.5 |  | 61.1 | 3.0 | 563 | 1.2 |
| 56.1 | 0.2 | 44 | 2.8 |  | 63.0 | 2.8 | 526 | 2.7 |
| 57.4 | 0.3 | 61 | 3.6 |  | 65.8 | 3.2 | 598 | 1.9 |
| 60.1 | 0.3 | 52 | 1.3 |  | 67.2 | 1.9 | 348 | 0.8 |
| 63.3 | 0.3 | 55 | 1.2 |  | 69.4 | 2.5 | 468 | 2.6 |
| 63.8 | 0.3 | 58 | 1.4 |  | 72.9 | 2.5 | 472 | 2.4 |
| 65.8 | 0.2 | 41 | 0.3 |  | 76.1 | 2.5 | 464 | 1.5 |
| 68.2 | 0.2 | 33 | 0.3 |  | 79.4 | 1.5 | 281 | 1.1 |
| 68.8 | 0.3 | 61 | 2.9 |  | 80.9 | 1.8 | 338 | 1.1 |
| 69.5 | 0.4 | 79 | 3.7 |  | 84.0 | 2.5 | 465 | 1.3 |
| 71.9 | 0.2 | 37 | 0.6 |  | 85.2 | 2.5 | 463 | 0.9 |
| 72.4 | 0.2 | 38 | 0.9 |  | 87.5 | 2.5 | 473 | 0.4 |
| 72.8 | 0.2 | 39 | 3.7 |  | 89.1 | 2.5 | 465 | 0.4 |
| 73.3 | 0.3 | 51 | 1.0 |  | 103.2 | 1.5 | 281 | 0.2 |
| 73.9 | 0.3 | 59 | 1.5 |  | 110.3 | 1.3 | 248 | 0.5 |
| 74.5 | 0.6 | 113 | 0.9 |  | 121.5 | 2.5 | 474 | 0.6 |
| 74.9 | 0.2 | 29 | 1.0 |  | 123.0 | 1.7 | 319 | 0.2 |
| 75.4 | 0.3 | 51 | 0.6 |  | 124.7 | 2.4 | 458 | 0.6 |
| 76.0 | 0.3 | 58 | 0.3 |  | 126.9 | 2.2 | 405 | 0.2 |
| 76.9 | 0.3 | 59 | 0.6 |  | 129.4 | 1.7 | 322 | 0.2 |
| 77.5 | 0.3 | 52 | 0.4 |  | 130.6 | 1.2 | 226 | 0.4 |
| 78.3 | 0.3 | 54 | 0.7 |  | 139.0 | 1.8 | 341 | 0.8 |
| 79.5 | 0.2 | 38 | 3.9 |  | 173.0 | 1.9 | 361 | 0.8 |
| 83.8 | 0.3 | 58 | 0.2 |  | 175.2 | 3.0 | 563 | 3.3 |
| 84.7 | 0.3 | 45 | - |  | 178.1 | 3.3 | 626 | 4.1 |
| 85.7 | 0.2 | 53 | 1.6 |  | 181.0 | 3.0 | 571 | 1.0 |
| 88.5 | 0.3 | 39 | 3.9 |  | 184.1 | 1.7 | 324 | 0.4 |
| 89.7 | 0.2 | 34 | 0.5 |  |  |  |  |  |
| 91.7 | 0.2 | 51 | 0.7 |  |  |  |  |  |
| 103.3 | 0.3 | 54 | 1.2 |  |  |  |  |  |
| 104.8 | 0.3 | 44 | 0.4 |  |  |  |  |  |
| 108.4 | 0.2 | 47 | 0.7 |  |  |  |  |  |
| 109.7 | 0.3 | 36 | 0.8 |  |  |  |  |  |
| 110.2 | 0.2 | 73 | 4.6 |  |  |  |  |  |
| 132.1 | 0.4 | 61 | 2.9 |  |  |  |  |  |

The comparison of the 1D ^13^C INEPT and CP spectra of *M. smegmatis* (blue) and *M. abscessus* (red) is shown in **Figure 2C**. To observe the relative differences in peak intensities, first we normalized the intensities of both spectra to the resonance at ∼32.5 ppm originating from mycolic acid signal, **Figure 2C top/middle**.^49^ The relative intensity of the INEPT signals in *M. smegmatis* is less compared to *M. abscessus* in particular in the arabinogalactan and peptidoglycan regions. On the other hand, the relative intensity of the CP signals in *M. smegmatis* is larger, in the arabinogalactan and peptidoglycan regions. These two comparisons indicate that *M. smegmatis* has relatively larger amount of rigid and smaller amount of flexible resonances compared to *M. abscessus*. In contrast, *M. abscessus* has relatively smaller amount of rigid and larger amount of flexible arabinogalactan and peptidoglycan resonances compared to *M. smegmatis*.

As an alternative method of comparing the relative abundance of PG, AG, MA signals for *M. smegmatis* and *M. abscessus*, we normalized the CP spectra at the CO region that is predominantly composed of PG signals, **Figure 2C bottom**. In this case, the signals originating from peptidoglycan and arabinogalactan have very similar intensities (so abundance) for both samples, and the predominant difference appears to be in the mycolic acid region, where *M. abscessus* has more mycolic acid abundance. In a recent work, The relative signal intensities in ^13^C CP ssNMR spectra were compared for these two bacteria prepared slightly differently by others,^49^ where the ^13^C spectra were shown to have intensity differences in the peptidoglycan region compared to our spectra. This could be due to different sample preparation, as well as different protocol used for normalization of the signals which has a large effect on the relative-abundance comparison. The comparison they made for these two NTM strains indicate similar behavior to our current results (with the MA normalization) where the PG signals were more abundant for *M. smegmatis*.

The CP buildup curves for *M. smegmatis* and *M. abscessus* are shown in **Figure 2D,E**, for four different chemical shift ranges in the ^13^C CP ssNMR spectra as depicted; 190-160, 160-110, 110-47 and 47-10 ppm. For a tentative group assignment of these resonances see **Figure 4**. The general behavior of the CP buildup curves for both samples are similar, indicating the similarity of the dynamics of the samples. We utilized a CP time of 1ms in recording the optimum ^13^C CPMAS spectra which allows a quantitative analysis of signal intensities. Moreover, these curves for NTM resemble the *Pseudomonas* biofilm CP buildup curves,^39^ and other polysaccharide and whole cell bacteria work,^55, 56^ where the maxima is observed at ∼500-1000 μs except for the carbonyl signals. The chemical shift region between 110-47 ppm (due to polysaccharides in PG, AG as well as peptide signals from PG) surprisingly shows a continuous decay towards longer buildup times. This decay is indicative of a rather rigid chemical site with different proton T_1p_ relaxation time that is dynamically different from other three resonance groups.

**Figure 4.**
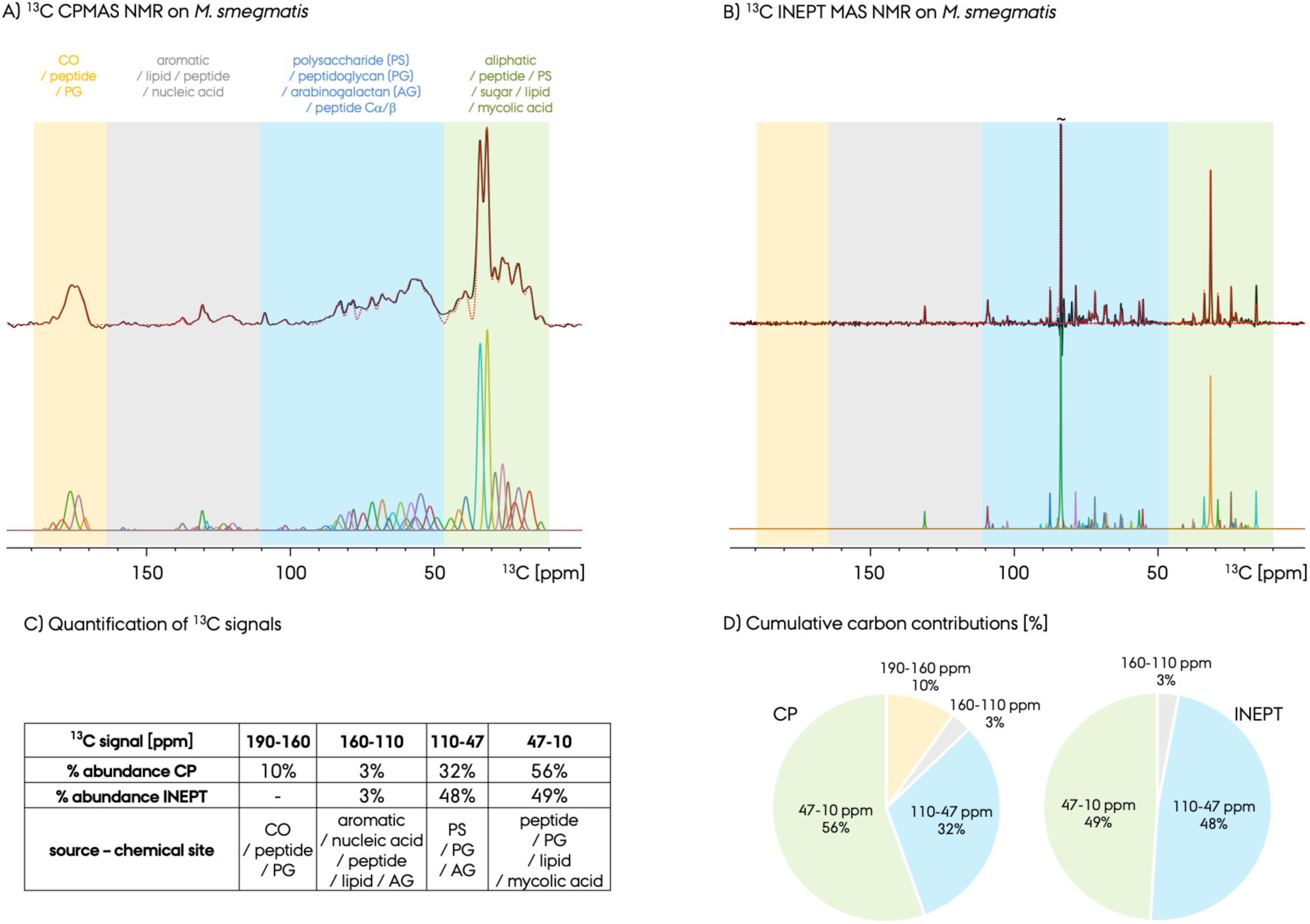
Quantification of different resonances in the fixed *M. smegmatis* by 1D ^13^C MAS ssNMR spectra recorded with **A**) CP and **B**) INEPT polarization transfer and peak deconvolution. The peaks are determined and fitted by ssNake program. Different chemical shift ranges are color coded and labeled in A. The bulk integration and corresponding percentages of chemical shift ranges are given in **C**) and **D**) for CP and INEPT spectra. The list of the identified cross peaks along with their abundance and linewidths are given below in Table 1.

We compared the ^13^C ssNMR spectra of *M. smegmatis* where we prepared the same batch of these sample with and without the fixation agent paraformaldehyde in PBS buffer, **Figure 3A**. We used the fixation protocol to preserve the freshly prepared bacterial samples and prevent changes or decay over time. The question arises with this treatment is weather the fixation cause changes in the samples, but the fixed versus unfixed (native) samples had the same NMR spectra. The overlay of the 1D ^13^C INEPT and INEPT-based 2D ^1^H-^13^C ssNMR spectra compare these two preparations, **Figure 3B,C**. The spectra overlap to a great extent which indicates that the fixation procedure has minimal effect on the sample.

The deconvolution analysis and tentative group assignments of the carbon chemical shifts observed in the CP and INEPT ^13^C ssNMR spectra are shown in **Figure 4A,B**. Previous bacterial cell wall studies and the CCMRD database on polysaccharide/carbohydrate chemical shifts were utilized to tentatively assign the peaks.^31, 39, 57, 58^ In this database, the polysaccharides are from various organisms, including bacteria, plants, fungi and represents the so far identified chemical sites and does not have complete coverage. Despite the differences in the exact chemical structure of these species and sparse coverage of the database, we utilized CCMRD hits for observed chemical shifts in the 2D ^1^H-^13^C spectrum to identify polysaccharides in our bacteria due to the overall similarities among species and their polysaccharides.^34^ The peak deconvolution analysis was done by utilizing ssNake fitting program package,^52^ a good agreement was achieved between the fit spectrum (dashed red) and the recorded one (solid black). We and others also recently showed the extent of information that can be obtained from overlapped 1D ^13^C spectra recorded with CP or INEPT, e.g. for *Pseudomonas* biofilm.^31, 39, 57^ The corresponding semi-quantitative abundance ratio analysis of NMR signals from different chemical shift regions are given in **Figure 4C,D**. The complete list of the identified chemical sites of ∼100 are listed in **Table 1**. Overall, the PG and peptide carbonyl signals are at ∼190 – 160 ppm; aromatic, peptide and lipid signals are at ∼160 – 110 ppm; polysaccharides, peptidoglycan, arabinogalactan and peptide (in PG or others) signals are at ∼110 – 47 ppm; and finally, the aliphatic signals from peptidoglycan, peptides, mycolic acid and lipids are at ∼47-10 ppm.

### 3.3. 2D ^1^H-^13^C MAS ssNMR spectroscopy for high-resolution native NTM characterization

Figure 5A-C depicts 2D ^1^H-^13^C ssNMR correlation spectra for native *M. smegmatis* and *M. abscessus* samples. The 2D spectra were recorded as carbon-detected INEPT-based HC and proton-detected INEPT/CP-based hCH spectra at 20 kHz MAS frequency and 275 K set temperature. The 2D carbon-detected CP-based HC on native *M. smegmatis* is shown in **Supplementary Figure 1**. The 2D spectra were recorded in ∼22 and ∼3-9 hours for the carbon-detected and proton-detected ssNMR experiments, respectively. The INEPT and CP based spectra selectively probes flexible versus rigid fraction of the signals from the samples, with significantly different resonance linewidths. Due to the detection of the flexible signals in the INEPT based spectra, the resolution is remarkably high with well resolved resonances observed allowing us to do tentative assignments. The CP based spectra are, however, very broad and only the strongest aliphatic region is observed due to the broader resonances from the rigid fraction of the sample, nevertheless overlapping well with the INEPT-based spectra.

**Figure 5.**
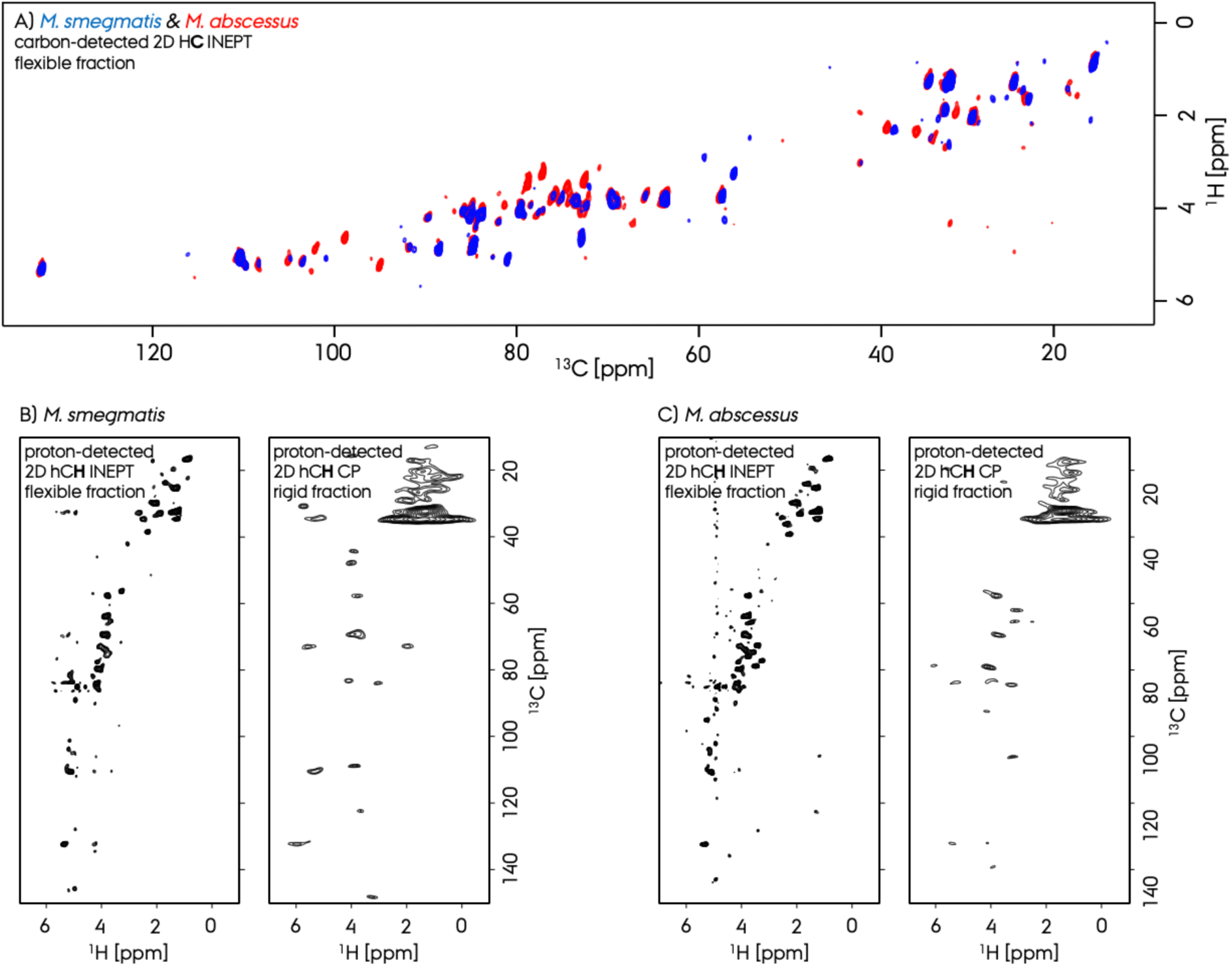
A) High-resolution 2D ^1^H-^13^C ssNMR correlation spectra for *M. smegmatis* and *M. abscessus* samples. The 2D spectra were recorded as carbon-detected INEPT-based HC and proton-detected INEPT/CP-based hCH spectra. The proton-detected 2D ^1^H-^13^C hCH ssNMR spectra for **B**) *M. smegmatis* and **C**) *M. abscessus*. at 20 kHz MAS frequency, 275 K set temperature and 750 MHz spectrometer by using a 3.2 mm thin-walled rotor. See experimental section for more details. The INEPT and CP based spectra selectively probes flexible versus rigid fraction of the signals.

Compared to the 1D ssNMR based chemical composition analysis described above, the INEPT based 2D spectra has high-resolution and provide a better means of identification of chemical sites, similar to our previous ssNMR study on a *Pseudomonas* biofilm.^39^ Similarly, we showed that the high-resolution 2D spectra on a *Pseudomonas* biofilm sample is indispensable to utilize for signal identification. In this work, due to the nature of the biofilm that is composed of polysaccharides in PG as well as secreted proteins, the protein specific signals were highly abundant, as abundant as the polysaccharide signals. However, the spectra of the NTM cell-envelope does not have significant protein and aromatic signals, it is instead mostly composed of signals from different types of polysaccharides, PG, AG, and mycolic acid.^49^ The comparison of the 2D carbon-detected INEPT-based HC spectra on *M. smegmatis*, and *Pseudomonas* biofilm is shown in **Supplementary Figure 2**. Nevertheless, we still observed chemical shifts specific to the peptide signals from the peptidoglycans specific to glycine, alanine, lysine and glutamine, Figure 6A. More details on the assignment of the polysaccharide resonances are given below in the next section.

**Figure 6.**
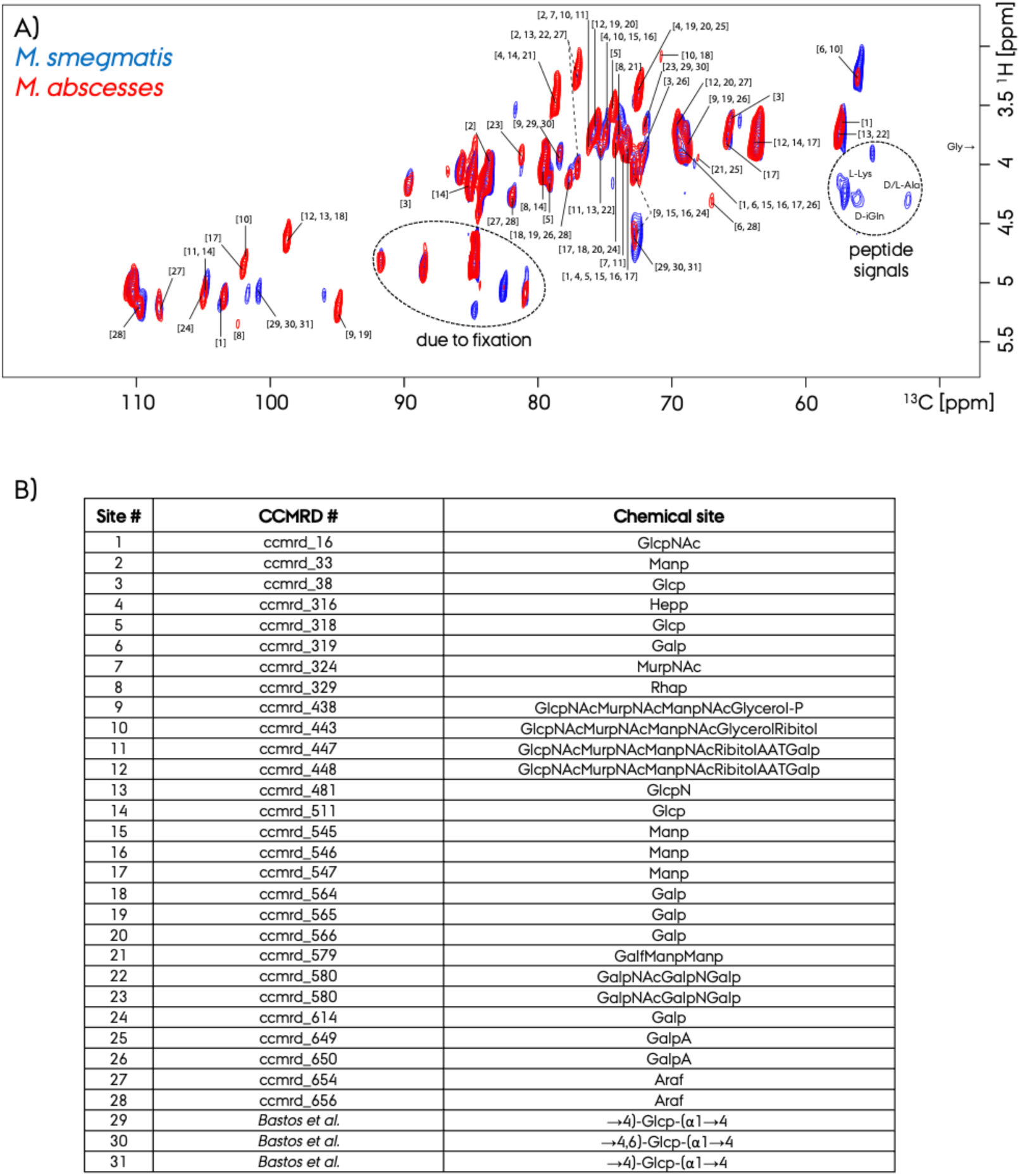
Representation of the tentative polysaccharide ^1^H/^13^C resonance assignments in the INEPT-based 2D ^1^H-^13^C ssNMR spectrum for *M. smegmatis* and *M. abscessus*. We utilized the chemical shift values and the CCMRD database for identification of species. We only judge the presence of a certain polysaccharide species with three to six cross peak pairs identified from a specific candidate. The complete list of the identified resonances are given in **Supplementary Table 1**. The peptide signals from PG are additionally highlighted.

The low-resolution (also lower sensitivity) CP-based 2D spectra is difficult to analyze and requires further advanced ssNMR instrumental tools, such as ultra-fast MAS of >100 kHz to get resolved and be helpful for resonance assignment.^59^ This approach would both increase the resolution and sensitivity of the proton-detected ssNMR spectroscopy. For this reason, we did not attempt to assign resonances in the CP-based 2D spectrum of the rigid NTM cell envelope fraction due to low resolution. Future studies utilizing MAS at >100 kHz could increase the sensitivity and resolution of the rigid fraction of NTM samples, as shown for bacterial cell wall peptidoglycan system.^46^ Alternative ssNMR approach such as hyperpolarized DNP ssNMR could increase the sensitivity by several orders of magnitude and allow 2D ^13^C-^13^C ssNMR spectra possible to record at natural abundance. A downside of DNP ssNMR is the lowering of the resolution due to cryogenic experimental temperatures. Fungal systems have been studied with such DNP ssNMR recently successfully.^60^ For the 2D ^13^C-^13^C ssNMR spectroscopy, ^13^C/^15^N isotope labeling will be very helpful for *in-vitro* prepared samples,^56, 57^ however, is not suitable for studying native materials such as the samples in this study or patient derived samples.

### 3.4. Polysaccharide compositional analysis of native NTM

By utilizing the high-resolution INEPT-based 2D ^1^H-^13^C ssNMR spectra we tentatively assigned resonances for the polysaccharides from the PG and AG fractions of the NTM cell envelope, Figure 6. CCMRD database was used for identifying the chemical shifts of resonances obtained in our spectra for the *M. smegmatis* and *M. abscessus*. We demonstrated this approach for a *Pseudomonas* biofilm recently and identified the major polysaccharide species present.^39^ In ssNMR based work, studies of complex native materials, such as the current NTM strains, missing peaks are common and the complete resonance set for a chemical species sometimes might not be observed due to the signal overlap or other experimental reasons,. For this reason, we judged the existence of a chemical species and assignment, with at least three identified resonances from that particular molecule to allow a reliable identification.

By following this principle, using the CCMRD database as well previous work on complex systems containing similar chemicals as NTM cell envelope, **31** unique polysaccharide signals were identified in the 2D spectra for the first time. These are glucose (Glcp), glucosamine (GlcpN), mannan (Manp), heptose (Hepp), galactose (Galp), galactoseamine (GalpN), N-acylated muramic acid (MurpNAc), N-acylated Glcp (GlcpNAc), galactomannan (GalfManp), galacturonic acid (GalpA) from the PG fraction, and arabinan (Araf) galactofuranose (Galf) and rhamnan (Rhap) from the AG fraction of the NTM cell envelope. The complete list of the identified resonances are given in **Supplementary Table 1**. Overall, our results are consistent with described polysaccharide components of the mycobacterial envelope.^16, 19^ These chemical species have not been identified previously in ssNMR studies for NTM due to the use of lower resolution 1D ^13^C ssNMR spectra on dried samples. NTM was characterized by identifying three main bulk chemical groups, peptidoglycan, arabinogalactan and mycolic acid, as the constituents of the NTM cell envelope.^49^

However, so far the identification of individual sugar types for NTM and also for many other systems have not been demonstrated by ssNMR. Our results highlight the vibrant nature of these NTM cell envelope components in their native environment. Current in depth chemical profiling of peptidoglycan, arabinogalactan and mycolic acid will open up novel sites to target against the fight with NTM bacteria. Remarkably, we identified several polysaccharide signals specific to either *M. smegmatis* or *M. abscessus*, Figure 6A, which indicate differences between these two bacteria.

## 4. Conclusions

In this work we present a high-resolution 1D and 2D ssNMR study on two NTM strains, *M. smegmatis* and *M. abscessus*. 1D and 2D MAS ^13^C-detected ssNMR spectra utilized here adds valuable structural and dynamics information input to the current state of the art. Native hydrated NTM samples were studied at natural abundance without isotope-labelling and any chemical or physical modification. We utilized 1D ^13^C and 2D ^1^H-^13^C ssNMR spectra and peak deconvolution to identify NTM cell envelope chemical sites. More than ∼100 distinct ^13^C signals were identified in the ssNMR spectra. The signals originating from both the flexible and rigid fractions of the native bacteria samples were selectively analyzed by utilizing either CP or INEPT based ^13^C ssNMR spectra. CP buildup curves provide insights into the dynamical similarity of the cell-wall components for NTM strains. Signals from peptidoglycan, arabinogalactan and mycolic acid were identified. We also provide tentative assignments for 31 polysaccharide species by using well resolved ^1^H/^13^C chemical shifts from the 2D INEPT-based ^1^H-^13^C ssNMR spectrum. As an orthogonal way of characterizing NTM, electron microscopy EM was used to provide spatial characterization. ssNMR and EM data suggest that *M. abscessus* cell envelope is composed of a smaller peptidoglycan layer which is more flexible compared to *M. smegmatis*, which may be related to its higher pathogenicity. Here in this work, we used of high-resolution 2D ssNMR first time to characterize native NTM strains and identified chemical sites. These results will aid the development of structure-based approaches to combat NTM infections.

## Data Availability

All data can be requested from the corresponding author.

## Acknowledgements

UA acknowledges financial support from University of Pittsburgh startup funding and the high-field NMR infrastructure at the Structural Biology Department, School of Medicine, University of Pittsburgh.

## Supplementary Information

**Supplementary Figure 1.**
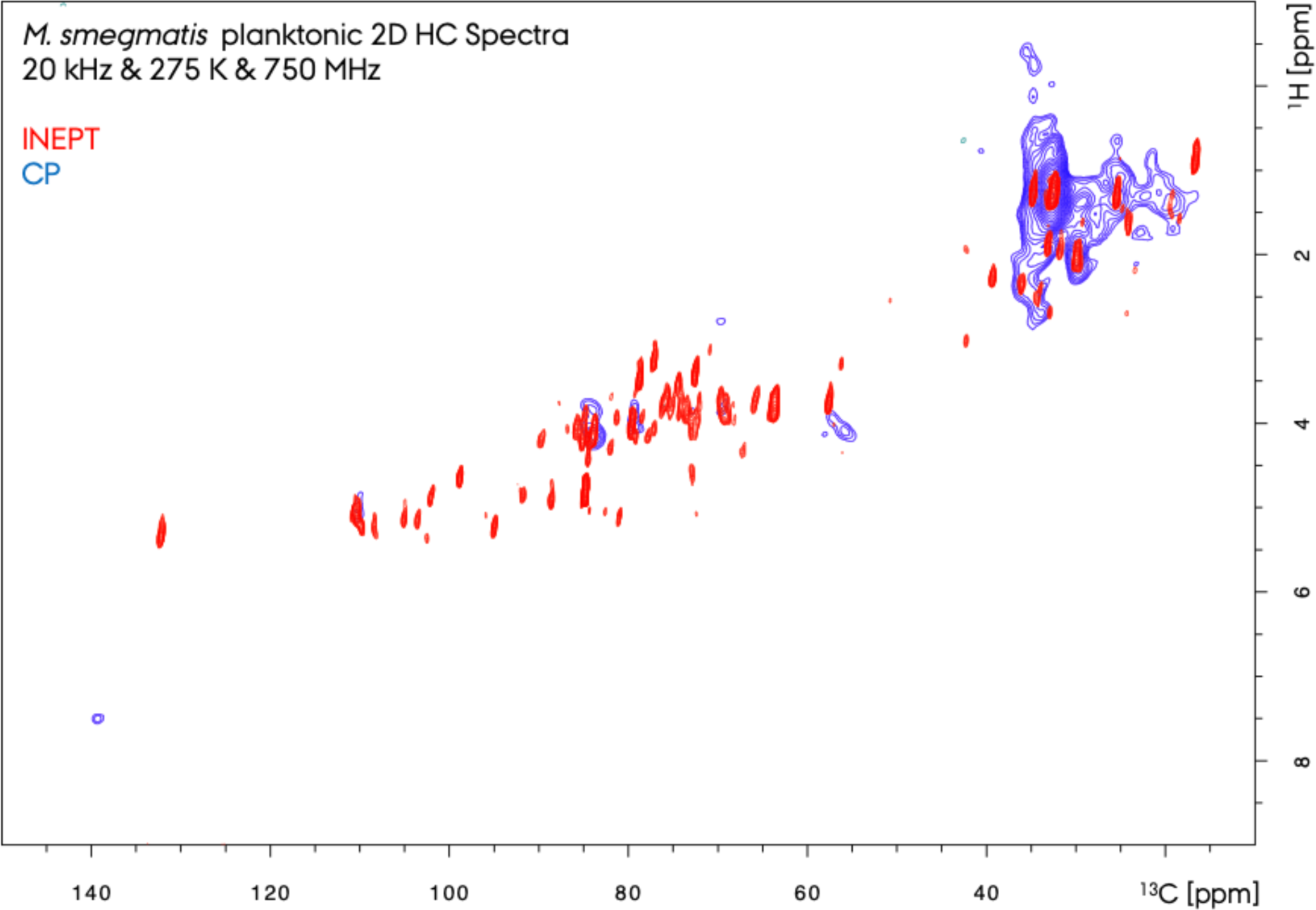
Comparison of the INEPT-based and CP-based 2D ^1^H-^13^C ssNMR spectrum for *M. smegmatis*.

**Supplementary Figure 2.**
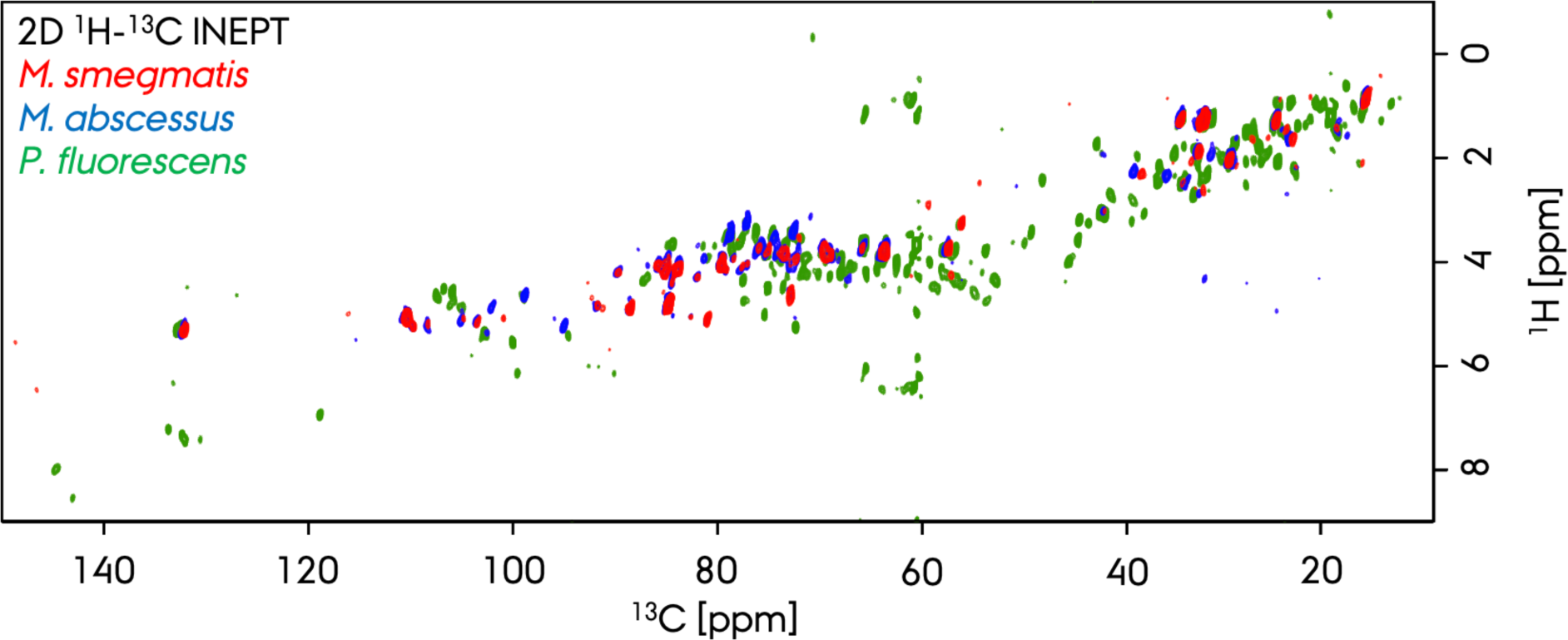
Comparison of the INEPT-based d 2D ^1^H-^13^C ssNMR spectra recorded for the *M. smegmatis*, *M. abscessus*. and *Pseudomonas fluorescens*.^39^

**Supplementary Table 1.**
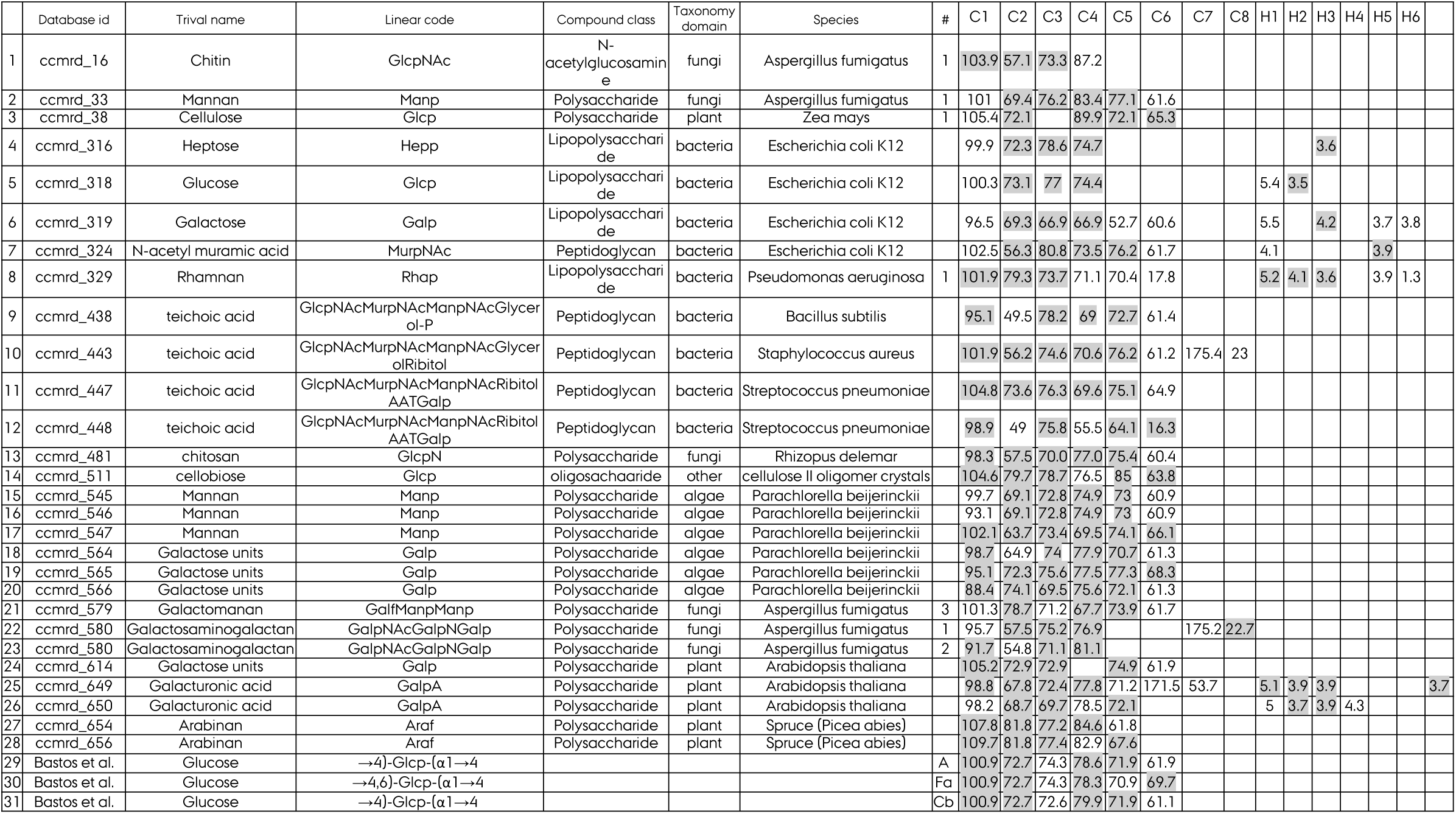
The list of the tentative polysaccharide ^1^H/^13^C resonance assignments in the INEPT-based 2D ^1^H-^13^C ssNMR spectrum for the *M. smegmatis* and *M. abscessus*. We utilized the chemical shift values and the CCMRD database for identification of species. We only judge the presence of a certain polysaccharide species with minimum three chemical shifts identified from a specific candidate. Entries 29-31 are not in the CCMRD, but from a publication.^48^

## References

(1) Walsh, T. R.; Gales, A. C.; Laxminarayan, R.; Dodd, P. C. Antimicrobial Resistance: Addressing a Global Threat to Humanity. PLoS Med 2023, 20 (7), e1004264. DOI: 10.1371/journal.pmed.1004264 From NLM.

(2) Wu, M. L.; Aziz, D. B.; Dartois, V.; Dick, T. NTM drug discovery: status, gaps and the way forward. Drug Discov Today 2018, 23 (8), 1502–1519. DOI: 10.1016/j.drudis.2018.04.001 From NLM.

(3) Kendall, B. A.; Varley, C. D.; Choi, D.; Cassidy, P. M.; Hedberg, K.; Ware, M. A.; Winthrop, K. L. Distinguishing tuberculosis from nontuberculous mycobacteria lung disease, Oregon, USA. Emerg Infect Dis 2011, 17 (3), 506–509. DOI: 10.3201/eid1703.101164 From NLM.

(4) Winthrop, K. L.; Marras, T. K.; Adjemian, J.; Zhang, H.; Wang, P.; Zhang, Q. Incidence and Prevalence of Nontuberculous Mycobacterial Lung Disease in a Large U.S. Managed Care Health Plan, 2008-2015. Ann Am Thorac Soc 2020, 17 (2), 178-185. DOI: 10.1513/AnnalsATS.201804-236OC From NLM.

(5) Johnson, M. M.; Odell, J. A. Nontuberculous mycobacterial pulmonary infections. J Thorac Dis 2014, 6 (3), 210–220. DOI: 10.3978/j.issn.2072-1439.2013.12.24 From NLM.

(6) Nohrenberg, M.; Wright, A.; Krause, V. Non-tuberculous mycobacterial skin and soft tissue infections in the Northern Territory, Australia, 1989-2021. Int J Infect Dis 2023, 135, 125-131. DOI: 10.1016/j.ijid.2023.07.031 From NLM.

(7) Máiz, L.; Girón, R.; Olveira, C.; Vendrell, M.; Nieto, R.; Martínez-García, M. A. Prevalence and factors associated with nontuberculous mycobacteria in non-cystic fibrosis bronchiectasis: a multicenter observational study. BMC Infect Dis 2016, 16 (1), 437. DOI: 10.1186/s12879-016-1774-x From NLM.

(8) Furukawa, B. S.; Flume, P. A. Nontuberculous Mycobacteria in Cystic Fibrosis. Semin Respir Crit Care Med 2018, 39 (3), 383–391. DOI: 10.1055/s-0038-1651495 From NLM.

(9) Floto, R. A.; Olivier, K. N.; Saiman, L.; Daley, C. L.; Herrmann, J. L.; Nick, J. A.; Noone, P. G.; Bilton, D.; Corris, P.; Gibson, R. L.;, et al. US Cystic Fibrosis Foundation and European Cystic Fibrosis Society consensus recommendations for the management of non-tuberculous mycobacteria in individuals with cystic fibrosis. Thorax 2016, 71 Suppl 1 (Suppl 1), i1-22. DOI: 10.1136/thoraxjnl-2015-207360 From NLM.

(10) Prevots, D. R.; Shaw, P. A.; Strickland, D.; Jackson, L. A.; Raebel, M. A.; Blosky, M. A.; Montes de Oca, R.; Shea, Y. R.; Seitz, A. E.; Holland, S. M.;, et al. Nontuberculous mycobacterial lung disease prevalence at four integrated health care delivery systems. Am J Respir Crit Care Med 2010, 182 (7), 970–976. DOI: 10.1164/rccm.201002-0310OC From NLM.

(11) Esther, C. R., Jr.; Esserman, D. A.; Gilligan, P.; Kerr, A.; Noone, P. G. Chronic Mycobacterium abscessus infection and lung function decline in cystic fibrosis. J Cyst Fibros 2010, 9 (2), 117–123. DOI: 10.1016/j.jcf.2009.12.001 From NLM.

(12) Qvist, T.; Taylor-Robinson, D.; Waldmann, E.; Olesen, H. V.; Hansen, C. R.; Mathiesen, I. H.; Høiby, N.; Katzenstein, T. L.; Smyth, R. L.; Diggle, P. J.;, et al. Comparing the harmful effects of nontuberculous mycobacteria and Gram negative bacteria on lung function in patients with cystic fibrosis. J Cyst Fibros 2016, 15 (3), 380–385. DOI: 10.1016/j.jcf.2015.09.007 From NLM.

(13) Lee, M. R.; Sheng, W. H.; Hung, C. C.; Yu, C. J.; Lee, L. N.; Hsueh, P. R. Mycobacterium abscessus Complex Infections in Humans. Emerg Infect Dis 2015, 21 (9), 1638–1646. DOI: 10.3201/2109.141634 From NLM.

(14) Koh, W. J.; Jeon, K.; Lee, N. Y.; Kim, B. J.; Kook, Y. H.; Lee, S. H.; Park, Y. K.; Kim, C. K.; Shin, S. J.; Huitt, G. A.;, et al. Clinical significance of differentiation of Mycobacterium massiliense from Mycobacterium abscessus. Am J Respir Crit Care Med 2011, 183 (3), 405–410. DOI: 10.1164/rccm.201003-0395OC From NLM.

(15) Philley, J. V.; DeGroote, M. A.; Honda, J. R.; Chan, M. M.; Kasperbauer, S.; Walter, N. D.; Chan, E. D. Treatment of Non-Tuberculous Mycobacterial Lung Disease. Curr Treat Options Infect Dis 2016, 8 (4), 275–296. DOI: 10.1007/s40506-016-0086-4 From NLM.

(16) Daffé, M.; Marrakchi, H. Unraveling the Structure of the Mycobacterial Envelope. Microbiol Spectr 2019, 7 (4). DOI: 10.1128/microbiolspec.GPP3-0027-2018 From NLM.

(17) Zuber, B.; Chami, M.; Houssin, C.; Dubochet, J.; Griffiths, G.; Daffé, M. Direct visualization of the outer membrane of mycobacteria and corynebacteria in their native state. J Bacteriol 2008, 190 (16), 5672–5680. DOI: 10.1128/jb.01919-07 From NLM.

(18) Hoffmann, C.; Leis, A.; Niederweis, M.; Plitzko, J. M.; Engelhardt, H. Disclosure of the mycobacterial outer membrane: cryo-electron tomography and vitreous sections reveal the lipid bilayer structure. Proc Natl Acad Sci U S A 2008, 105 (10), 3963–3967. DOI: 10.1073/pnas.0709530105 From NLM.

(19) Angala, S. K.; Palčeková, Z.; Belardinelli, J. M.; Jackson, M. Covalent modifications of polysaccharides in mycobacteria. Nat Chem Biol 2018, 14 (3), 193–198. DOI: 10.1038/nchembio.2571 From NLM.

(20) Batt, S. M.; Burke, C. E.; Moorey, A. R.; Besra, G. S. Antibiotics and resistance: the two-sided coin of the mycobacterial cell wall. Cell Surf 2020, 6, 100044. DOI: 10.1016/j.tcsw.2020.100044 From NLM.

(21) Ojha, A. K.; Baughn, A. D.; Sambandan, D.; Hsu, T.; Trivelli, X.; Guerardel, Y.; Alahari, A.; Kremer, L.; Jacobs, W. R.; Hatfull, G. F. Growth of Mycobacterium tuberculosis biofilms containing free mycolic acids and harbouring drug-tolerant bacteria. Molecular Microbiology 2008, 69 (1), 164–174. DOI: 10.1111/j.1365-2958.2008.06274.x.

(22) Wiersma, C. J.; Belardinelli, J. M.; Avanzi, C.; Angala, S. K.; Everall, I.; Angala, B.; Kendall, E.; de Moura, V. C. N.; Verma, D.; Benoit, J.;, et al. Cell Surface Remodeling of Mycobacterium abscessus under Cystic Fibrosis Airway Growth Conditions. ACS Infect Dis 2020, 6 (8), 2143–2154. DOI: 10.1021/acsinfecdis.0c00214 From NLM.

(23) Kalscheuer, R.; Palacios, A.; Anso, I.; Cifuente, J.; Anguita, J.; Jacobs, W. R., Jr.; Guerin, M. E.; Prados-Rosales, R. The Mycobacterium tuberculosis capsule: a cell structure with key implications in pathogenesis. Biochem J 2019, 476 (14), 1995–2016. DOI: 10.1042/bcj20190324 From NLM.

(24) Jarlier, V.; Nikaido, H. Permeability barrier to hydrophilic solutes in Mycobacterium chelonei. J Bacteriol 1990, 172 (3), 1418–1423. DOI: 10.1128/jb.172.3.1418-1423.1990 From NLM.

(25) Rhoades, E. R.; Archambault, A. S.; Greendyke, R.; Hsu, F. F.; Streeter, C.; Byrd, T. F. Mycobacterium abscessus Glycopeptidolipids mask underlying cell wall phosphatidyl-myo-inositol mannosides blocking induction of human macrophage TNF-alpha by preventing interaction with TLR2. J Immunol 2009, 183 (3), 1997–2007. DOI: 10.4049/jimmunol.0802181 From NLM.

(26) Tuttle, M. D.; Comellas, G.; Nieuwkoop, A. J.; Covell, D. J.; Berthold, D. A.; Kloepper, K. D.; Courtney, J. M.; Kim, J. K.; Barclay, A. M.; Kendall, A.;, et al. Solid-state NMR structure of a pathogenic fibril of full-length human alpha-synuclein. Nature Structural & Molecular Biology 2016, 23 (5), 409–415. DOI: 10.1038/nsmb.3194.

(27) Asano, S.; Engel, B. D.; Baumeister, W. In Situ Cryo-Electron Tomography: A Post-Reductionist Approach to Structural Biology. Journal of Molecular Biology 2016, 428 (2), 332–343. DOI: 10.1016/j.jmb.2015.09.030.

(28) Plitzko, J. M.; Schuler, B.; Selenko, P. Structural Biology outside the box - inside the cell. Current Opinion in Structural Biology 2017, 46, 110–121. DOI: 10.1016/j.sbi.2017.06.007.

(29) Earl, L. A.; Falconieri, V.; Subramaniam, S. Microbiology catches the cryo-EM bug. Current Opinion in Microbiology 2018, 43, 199–207. DOI: 10.1016/j.mib.2018.02.012.

(30) Narasimhan, S.; Folkers, G. E.; Baldus, M. When Small becomes Too Big: Expanding the Use of In-Cell Solid-State NMR Spectroscopy. Chempluschem 2020, 85 (4), 760–768. DOI: 10.1002/cplu.202000167.

(31) Romaniuk, J. A. H.; Cegelski, L. Bacterial cell wall composition and the influence of antibiotics by cell-wall and whole-cell NMR. Philosophical Transactions of the Royal Society B-Biological Sciences 2015, 370 (1679). DOI: 10.1098/rstb.2015.0024.

(32) Jeffries, J.; Thongsomboon, W.; Visser, J. A.; Enriquez, K.; Yager, D.; Cegelski, L. Variation in the ratio of curli and phosphoethanolamine cellulose associated with biofilm architecture and properties. Biopolymers 2021, 112 (1). DOI: 10.1002/bip.23395.

(33) Reichhardt, C.; Joubert, L. M.; Clemons, K. V.; Stevens, D. A.; Cegelski, L. Integration of electron microscopy and solid-state NMR analysis for new views and compositional parameters of Aspergillus fumigatus biofilms. Medical Mycology 2019, 57, S239–S244. DOI: 10.1093/mmy/myy140.

(34) Ghassemi, N.; Poulhazan, A.; Deligey, F.; Mentink-Vigier, F.; Marcotte, I.; Wang, T. Solid-State NMR Investigations of Extracellular Matrixes and Cell Walls of Algae, Bacteria, Fungi, and Plants. Chemical Reviews 2022, 122 (10), 10036–10086. DOI: 10.1021/acs.chemrev.1c00669.

(35) Manz, C.; Pagel, K. Glycan analysis by ion mobility-mass spectrometry and gas-phase spectroscopy. Curr Opin Chem Biol 2018, 42, 16–24. DOI: 10.1016/j.cbpa.2017.10.021 From NLM.

(36) Ndukwe, I. E.; Black, I.; Heiss, C.; Azadi, P. Evaluating the Utility of Permethylated Polysaccharide Solution NMR Data for Characterization of Insoluble Plant Cell Wall Polysaccharides. Analytical Chemistry 2020, 92 (19), 13221–13228. DOI: 10.1021/acs.analchem.0c02379.

(37) Poulhazan, A.; Widanage, M. C. D.; Muszynprimeski, A.; Arnold, A. A.; Warschawski, D. E.; Azadi, P.; Marcotte, I.; Wang, T. Identification and Quantification of Glycans in Whole Cells: Architecture of Microalgal Polysaccharides Described by Solid-State Nuclear Magnetic Resonance. Journal of the American Chemical Society 2021, 143 (46), 19374–19388. DOI: 10.1021/jacs.1c07429.

(38) Kang, X.; Kirui, A.; Muszynski, A.; Widanage, M. C. D.; Chen, A.; Azadi, P.; Wang, P.; Mentink-Vigier, F.; Wang, T. Molecular architecture of fungal cell walls revealed by solid-state NMR. Nature Communications 2018, 9. DOI: 10.1038/s41467-018-05199-0.

(39) Byeon, C.-H.; Kinney, T.; Saricayir, H.; Srinivasa, S.; Wells, M. K.; Kim, W.; Akbey, Ü. Tapping into the native Pseudomonas bacterial biofilm structure by high-resolution multidimensional solid-state NMR. Journal of Magnetic Resonance 2023, 357, 107587. DOI: 10.1016/j.jmr.2023.107587.

(40) Jennings, L. K.; Dreifus, J. E.; Reichhardt, C.; Storek, K. M.; Secor, P. R.; Wozniak, D. J.; Hisert, K. B.; Parsek, M. R. Pseudomonas aeruginosa aggregates in cystic fibrosis sputum produce exopolysaccharides that likely impede current therapies. Cell Reports 2021, 34 (8). DOI: 10.1016/j.celrep.2021.108782.

(41) Kirui, A.; Zhao, W. C.; Deligey, F.; Yang, H.; Kang, X.; Mentink-Vigier, F.; Wang, T. Carbohydrate-aromatic interface and molecular architecture of lignocellulose. Nature Communications 2022, 13 (1). DOI: 10.1038/s41467-022-28165-3.

(42) Kang, X.; Kirui, A.; Widanage, M. C. D.; Mentink-Vigier, F.; Cosgrove, D. J.; Wang, T. Lignin-polysaccharide interactions in plant secondary cell walls revealed by solid-state NMR. Nature Communications 2019, 10. DOI: 10.1038/s41467-018-08252-0.

(43) Takahashi, H.; Ayala, I.; Bardet, M.; De Paepe, G.; Simorre, J. P.; Hediger, S. Solid-State NMR on Bacterial Cells: Selective Cell Wall Signal Enhancement and Resolution Improvement using Dynamic Nuclear Polarization. Journal of the American Chemical Society 2013, 135 (13), 5105–5110. DOI: 10.1021/ja312501d.

(44) Nygaard, R.; Romaniuk, J. A. H.; Rice, D. M.; Cegelski, L. Spectral Snapshots of Bacterial Cell-Wall Composition and the Influence of Antibiotics by Whole-Cell NMR. Biophysical Journal 2015, 108 (6), 1380–1389. DOI: 10.1016/j.bpj.2015.01.037.

(45) Laguri, C.; Silipo, A.; Martorana, A. M.; Schanda, P.; Marchetti, R.; Polissi, A.; Molinaro, A.; Simorre, J. P. Solid State NMR Studies of Intact Lipopolysaccharide Endotoxin. Acs Chemical Biology 2018, 13 (8), 2106–2113. DOI: 10.1021/acschembio.8b00271.

(46) Bougault, C.; Ayala, I.; Vollmer, W.; Simorre, J. P.; Schanda, P. Studying intact bacterial peptidoglycan by proton-detected NMR spectroscopy at 100 kHz MAS frequency. Journal of Structural Biology 2019, 206 (1), 66–72. DOI: 10.1016/j.jsb.2018.07.009.

(47) Jean, N. L.; Bougault, C. M.; Egan, A. J. F.; Vollmer, W.; Simorre, J. P. Solution NMR assignment of LpoB, an outer-membrane anchored Penicillin-Binding Protein activator from *Escherichia coli*. Biomolecular Nmr Assignments 2015, 9 (1), 123–127. DOI: 10.1007/s12104-014-9557-z.

(48) Bastos, R.; Marín-Montesinos, I.; Ferreira, S. S.; Mentink-Vigier, F.; Sardo, M.; Mafra, L.; Coimbra, M. A.; Coelho, E. Covalent connectivity of glycogen in brewer’s spent yeast cell walls revealed by enzymatic approaches and dynamic nuclear polarization NMR. Carbohydrate Polymers 2024, 324, 121475. DOI: 10.1016/j.carbpol.2023.121475.

(49) Liu, X.; Brčić, J.; Cassell, G. H.; Cegelski, L. CPMAS NMR platform for direct compositional analysis of mycobacterial cell-wall complexes and whole cells. Journal of Magnetic Resonance Open 2023, 16–17, 100127. DOI: 10.1016/j.jmro.2023.100127.

(50) DePas, W. H.; Bergkessel, M.; Newman, D. K. Aggregation of Nontuberculous Mycobacteria Is Regulated by Carbon-Nitrogen Balance. mBio 2019, 10 (4). DOI: 10.1128/mBio.01715-19 From NLM.

(51) Wishart, D. S.; Bigam, C. G.; Holm, A.; Hodges, R. S.; Sykes, B. D. H-1, C-13 AND N-15 RANDOM COIL NMR CHEMICAL-SHIFTS OF THE COMMON AMINO-ACIDS .1. INVESTIGATIONS OF NEAREST-NEIGHBOR EFFECTS. Journal of Biomolecular Nmr 1995, 5 (1), 67-81. DOI: 10.1007/bf00227471.

(52) van Meerten, S. G. J.; Franssen, W. M. J.; Kentgens, A. P. M. ssNake: A cross-platform open-source NMR data processing and fitting application. Journal of Magnetic Resonance 2019, 301, 56–66, Article. DOI: 10.1016/j.jmr.2019.02.006.

(53) Tong, G.; Pan, Y.; Dong, H.; Pryor, R.; Wilson, G. E.; Schaefer, J. Structure and dynamics of pentaglycyl bridges in the cell walls of Staphylococcus aureus by C-13-N-15 REDOR NMR. Biochemistry 1997, 36, 9859.

(54) Alderwick, L. J.; Harrison, J.; Lloyd, G. S.; Birch, H. L. The Mycobacterial Cell Wall--Peptidoglycan and Arabinogalactan. Cold Spring Harb Perspect Med 2015, 5 (8), a021113. DOI: 10.1101/cshperspect.a021113 From NLM.

(55) Reichhardt, C. The Pseudomonas aeruginosa Biofilm Matrix Protein CdrA Has Similarities to Other Fibrillar Adhesin Proteins. Journal of Bacteriology 2023, 205 (5). DOI: 10.1128/jb.00019-23.

(56) Romaniuk, J. A. H.; Cegelski, L. Peptidoglycan and Teichoic Acid Levels and Alterations in Staphylococcus aureus by Cell-Wall and Whole-Cell Nuclear Magnetic Resonance. Biochemistry 2018, 57 (26), 3966–3975. DOI: 10.1021/acs.biochem.8b00495.

(57) Patti, G. J.; Kim, S. J.; Schaefer, J. Characterization of the peptidoglycan of vancomycin-susceptible Enterococcus faecium. Biochemistry 2008, 47 (32), 8378–8385. DOI: 10.1021/bi8008032.

(58) Kang, X.; Zhao, W. C.; Widanage, M. D. C.; Kirui, A.; Ozdenvar, U.; Wang, T. CCMRD: a solid-state NMR database for complex carbohydrates. Journal of Biomolecular Nmr 2020, 74 (4-5), 239–245. DOI: 10.1007/s10858-020-00304-2.

(59) Le Marchand, T.; Schubeis, T.; Bonaccorsi, M.; Paluch, P.; Lalli, D.; Pell, A. J.; Andreas, L. B.; Jaudzems, K.; Stanek, J.; Pintacuda, G. 1H-Detected Biomolecular NMR under Fast Magic-Angle Spinning. Chemical Reviews 2022, 122 (10), 9943–10018. DOI: 10.1021/acs.chemrev.1c00918.

(60) Fernando, L. D.; Widanage, M. C. D.; Shekar, S. C.; Mentink-Vigier, F.; Wang, P.; Wi, S.; Wang, T. Solid-state NMR analysis of unlabeled fungal cell walls from Aspergillus and Candida species. Journal of Structural Biology-X 2022, 6. DOI: 10.1016/j.yjsbx.2022.100070.

